# Multiple stressors and recruitment failure of long-lived endangered freshwater mussels with a complex life cycle

**DOI:** 10.1101/2021.09.20.461153

**Authors:** Kazuki Miura, Nobuo Ishiyama, Junjiro N. Negishi, Daisetsu Ito, Keita Kawajiri, Hokuto Izumi, Takahiro Inoue, Masahiro Nakaoka, Futoshi Nakamura

## Abstract

1. Multiple stressors can interactively affect the population of organisms; however, the process by which they affect recruitment efficiency remains unclear for empirical populations. Recruitment efficiency can be regulated at multiple stages of life, particularly in organisms with complex life cycles. Understanding the interactive effects of multiple stressors on recruitment efficiency and determining the bottleneck life stages is imperative for species conservation.
2. The proportion of <20-year-old juveniles of the endangered freshwater pearl mussel *Margaritifera togakushiensis*, which has an obligate parasitic larval stage, was investigated in 24 rivers from eastern Hokkaido, northern Japan to reveal the influence of nutrients, fine sediment, and their combined effects on juvenile recruitment efficiency. The following indices for recruitment at adult, parasitic, and post-parasitic juvenile stages were obtained from 11 of these rivers: gravid female density, glochidia density (the number of glochidia infections per stream area), and juvenile survival rate. This study explored the bottleneck stages of recruitment efficiency and the interactive effects of the two stressors on these stages.
3. Twenty-four population status assessments determined that the proportion of juveniles ranged from 0.00 to 0.53, and juveniles were absent from four rivers. The results showed that the parasitic and post-parasitic juvenile stages were bottlenecks for recruitment efficiency. Juvenile survival rates had a more significant positive effect on recruitment efficiency in rivers with a high glochidia density. Juvenile survival rate was decreased by the synergistic interaction of nutrients and fine sediment, although factors limiting glochidia density were not found.
4. The nutrient concentration of rivers in the study region was well explained by the proportion of agricultural land cover and urban areas in the watersheds, but no relationship was detected between fine sediment abundance and land use.
5. This study suggests that nutrient management at a catchment scale can be effective for re-establishing the recruitment of *M. togakushiensis*, particularly in rivers with a high content of fine sediments. The results also emphasise the importance of considering both parasitic and post-parasitic juvenile stages of mussels to maximise the positive effects of stressor mitigation.

## Introduction

Anthropogenic stressors have diminished the populations of various organisms worldwide (Hof et al., 2011; Reid et al., 2018). Many types of ecosystems are interactively affected by multiple stressors, which makes it difficult to predict the responses of organisms with a single factor (Darling & Côté, 2008). However, this highlights the importance of efforts to understand causal mechanisms for conservation biology (Hof et al., 2011). Studies on multiple stressors have mostly focussed on abundance or presence/absence data (e.g., Toft et al., 2018). Recruitment failure is also an important process in population decline (Strayer & Malcom, 2012). Low recruitment efficiencies can severely increase the risks of local extinctions, but there remain opportunities for recovering declined populations in situations where reproducing individuals are still present (Hylander & Ehrlén, 2013). However, few studies have demonstrated the interactive effects of stressors on recruitment efficiency.

The recruitment efficiency of organisms can be regulated at multiple stages of life. In particular, organisms with complex life cycles are sensitive to human disturbances (Kingsolver et al., 2011). These organisms have distinct life stages and an abrupt ontogenetic change in an individual’s morphology, physiology, and behaviour, which is usually associated with a change in habitat (Wilbur, 1980). This is because each life stage may contribute differently to recruitment efficiency and would be constrained by different stressors (González-Varo et al. 2012). To effectively conserve organisms with complex life cycles, it is crucial to identify the life stages that experience bottlenecks in recruitment efficiency by considering the interactive effects between multiple stressors. However, such empirical studies are very limited.

Freshwater mussels (Order Unionoida) are one of the most endangered organisms (Lopes-Lima et al., 2018). Their vulnerability is largely derived from their complex life cycles, which require host fish parasitisation during their larval phase (Haag, 2012). Many mussel populations face a high risk of extinction due to recruitment failure (Strayer & Malcom, 2012), especially if recruitment does not recover (Haag, 2012). Their longevity (4–190 years) allows them to persist for a long time, even in non-recruiting populations (Geist, 2010; Haag & Rypel, 2011). Recruitment should be urgently re-established before the disappearance of reproducing mussel individuals and timely intervention to prevent the local extinction of mussel populations.

Previous studies suggest that habitat degradation is a major cause of recruitment failure in freshwater mussels, and the benthic juveniles in their post-parasitic stage are considered to be a critical stage in their life cycle (Haag et al. 2019; Österling et al. 2010; Österling et al. 2008; Strayer & Malcom, 2012). Most of these studies have focussed on only part of their life cycle, which does not provide sufficient information for recovering their natural recruitment with actual conservation measures. This is because impaired recruitment might have been caused by problems at multiple stages of their life cycles (Galbraith et al., 2018). A comprehensive test is necessary to determine the life stage that regulates recruitment and the mechanisms and drivers that cause recruitment failure. Increased nutrients and fine sediment delivered from watersheds are considered to be major stressors threatening freshwater mussels worldwide (Lopes-Lima et al., 2017). In particular, benthic juveniles of lotic mussel species are sensitive to eutrophication and siltation because they require well-oxygenated substrates (Geist & Auerswald, 2007).

The present study aimed to reveal: 1) the key life stages limiting recruitment efficiency of freshwater pearl mussel *Margaritifera togakushiensis* (“FPM” hereafter); and 2) the relationship between the abundance of nutrients and fine sediment and their combined effects on recruitment failure. FPM is distributed in cold-water rivers in Sakhalin, Russia, and the Honshu and Hokkaido Islands of Japan (Miura, Ishiyama, Kawajiri, Atsumi, Nagasaka, et al., 2019). This species has recently been threatened by anthropogenic disturbances, such as habitat fragmentation, river habitat modification, and land-use change (Ministry of the Environment of Japan, 2014). First, we summarised the general life cycles of freshwater mussels and explained the reproduction processes and related important indices of their recruitment. In addition, this summary explained how to demonstrate the key life stages limiting their recruitment efficiency. Second, we determined the population status of FPM in 24 rivers in northern Japan. Third, we identified the key life stages using field observations and experiments. Finally, we explored the impacts of nutrients and fine sediment and their combined effects on the key life stages of FPM.

## Theoretical framework

### Adult stage

Adult freshwater mussels, including FPM, are typically sedentary, with limited dispersal ability (Appendix 1: Fig. S1) (Geist, 2010). The FPM has a shell length of <100 mm, and its reproductive season is from March to May (Kondo, 2008; Kurihara & Goto, 2011). They are generally gonochoristic. Mature females inhale sperm spheres released from mature males, and the fertilised eggs produce glochidium larvae in their gills (Haag, 2012). Mature glochidia are then released into the water column by the females. Therefore, the maximum larval production of the population is limited by the abundance of mature gravid individuals, and adequate larval production is necessary for high recruitment efficiency (McLain & Ross, 2005). Gravidity rate (the proportion of gravid individuals within the population) has been used as an index of fertilisation success and the production of mature glochidium larvae (Österling et al., 2008). The gravid abundance of the population can be described as the product of the gravidity rate and adult mussel density; this is defined as the “gravid density,” and is used as an index of the potential amount of larval production in this study.

### Parasitic stage

Glochidia released from female mussels attach to the gill tissue of a host fish and proceed to their parasitic stage (Appendix 1: Fig. S1) (Haag 2012). FPM mainly uses white-spotted char *Salvelinus leucomaenis leucomaenis* and perhaps Dolly Varden char *S. malma krascheninnikovi* as their host (Kondo, 2008; Kurihara & Goto, 2011). Those attached glochidia that fail to encyst or resist host immunity perish within several days to two weeks (Ziuganov et al. 1994). Therefore, the maximum number of recruited juveniles is limited by glochidia surviving this initial parasitisation period. The infection success of glochidia can be judged by observing the host tissues two weeks after attachment (Österling et al., 2008). Encysted glochidia live on host fish for several days to several months (Haag, 2012). FPM parasitises the gills of host fish for 40–50 days (Kondo, 2008). After metamorphosis, juveniles detach themselves from the host fish and reach the riverbed. A sufficient number of juveniles entering the population are necessary for high recruitment efficiency (Österling et al., 2008). The abundance of recruiting juveniles can be described as the product of host fish density, and the mean glochidia load (mean number of glochidia infections/fish in a population) on a host fish is referred to as “glochidia density” (Österling et al., 2008).

### Post-parasitic juvenile stage

The surviving juveniles proceeded to the post-parasitic juvenile stage (Appendix 1: Fig. S1). Post-parasitic juveniles live in riverbed sediments for several years to decades (Geist, 2010). The survival rate of juveniles directly affects the recruitment efficiency of freshwater mussels. This early juvenile stage is generally considered as the life stage that is most sensitive to habitat degradation (Buddensiek, 1995). Thus, this study used the “juvenile survival rate” shortly after the parasitic period as an index for this bottleneck stage.

### An empirical testing approach

Recruitment of freshwater mussels can be slowed or stopped (i.e. decreasing the proportion of juveniles or absence of juveniles) when any stage of their life cycle is disrupted (Österling et al., 2008). Therefore, the key life-cycle stage(s) regulating recruitment efficiency can be identified by examining the positive relationships between the degree of successful recruitment and the index of each life stage (i.e. gravid density, glochidia density, and juvenile survival rate) (Stoeckl et al. 2015). This study used the proportion of <20-year-old FPM individuals within the population (“the proportion of juveniles” hereafter) as an index for the degree of successful recruitment.

## Materials and methods

### Study site

The study was conducted in 24 rivers (sites) across eight watersheds encompassing >1500 km^2^ of eastern Hokkaido, Japan, where several populations of FPM were confirmed by Miura, Ishiyama, Kawajiri, Atsumi, Nagasaka, et al. (2019) (Fig. 1). Sites *a* to *o* and *p* to *x* run-off into Nemuro Strait and the Pacific Ocean, respectively. No sites were located upstream of other sites. The detailed locations of the sites remain undisclosed to assist the conservation of the target species. There were no large dams and weirs inhibiting fish migration. Another congeneric species, *Margaritifera laevis*, also occurs in the study region (Miura, Ishiyama, Kawajiri, Atsumi, Nagasaka, et al., 2019). The stone loach *Noemachelius barbatulus toni*, white-spotted char, and sticklebacks *Pungitius* spp. are the most dominant fish species in the studied streams (Ishiyama et al. 2020).

**Fig. 1.**
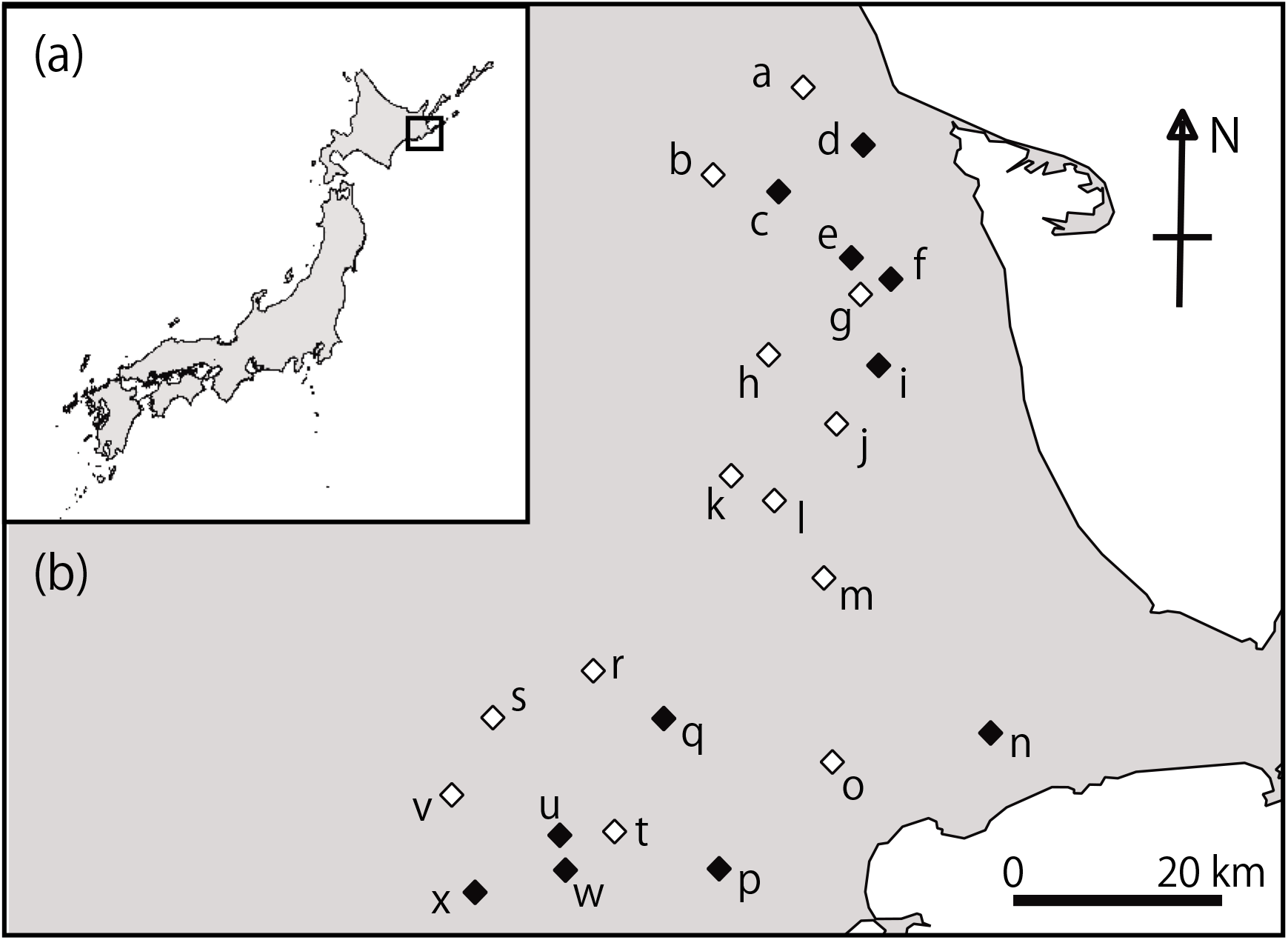
Geographical location of the 24 study rivers (sites) including 11 intensive study sites. White and black diamonds denote non-intensive and intensive sites, respectively.

### Population status assessment

Field sampling was carried out from June to September 2017 to assess the population status of FPM. Two study reaches [length: 26.9 ± 22.83 m (mean ± SD)] were established in glides at each site and were separated from each other by >500 m. Quadrat-based quantitative and semi-quantitative protocols were adopted to collect >50 individuals of FPM from each site (Appendix 2). Collected mussels were photographed for size measurements and released to the original study reach. The shell length and height of each individual were estimated using Image J software (Schneider et al. 2012) with species identification using non-lethal identification criteria (Miura, Ishiyama, Kawajiri, Atsumi, Tachibana, et al., 2019). The age of each individual was estimated using growth models (see Appendix 3). The histogram of the age composition of each population combined the data from the two reaches.

### Intensive field observations and experiments

Intensive field observations and experiments were conducted in one reach of 11 sites (Fig. 1) selected because they were easily accessible and had wide ranges in proportion to juveniles (0.00–0.53) and adult mussel density (1.85–40.00 individuals/m^2^) without correlations with each other (*r* = −0.13, *P* = 0.70).

The gravidity rate of FPM, which is a factor of the gravid density, was measured on 3– 6 May and 9–11 May 2018 (see “Appendix 4” for the method). These periods coincided with the peak of the FPM glochidia release (Kobayashi & Kondo 2009). Additional observations on 17– 18 May 2018 confirmed the approximate end of mussel gravidity and helped decide the timing of host fish sampling (see next paragraph). The adult density (N/m^2^) at each site was calculated using quadrat-based data obtained from field observations for population status assessment. The gravid density (N/m^2^) of each site was calculated by multiplying the gravidity rate with the adult density.

Electrofishing was performed using a backpack electrofisher (Model 12 B, Smith-Root Inc., Vancouver, WA, USA) between 6 June and 11 June 2018 to examine glochidia density, which is the parasitic period of FPM that occurs at least two weeks after their gravidity period in this region (Kurihara & Goto, 2011). At each site, one reach 10 times its mean wetted width was established near (upstream or downstream) the mussel monitoring area. Quantitative and qualitative electrofishing was conducted to collect host fish samples to calculate host density (N/m^2^) and mean glochidia load (see Appendix 5). Young-of-the-year (YOY) host fish are generally important for the recruitment of *Margaritifera* because they have not acquired immunity (Österling, 2015). However, preliminary observations showed encystment by host tissues and growth of glochidia on gills of ≥1^+^ host fish; therefore, both YOY and ≥1^+^ host individuals were collected and distinguished by the differences in fork length (mm) (Appendix 1: Fig. S2).

Glochidia density was calculated as follows: First, the mean glochidia load was calculated separately for the two age classes (YOY and ≥1^+^) because their body sizes differed greatly (Appendix 1: Fig. S2). Second, the host densities of each size class were multiplied by the mean glochidia load for each age class. Third, the glochidia density was calculated as the sum of the products of host density and mean glochidia load in each of the two age classes. When only one individual YOY was captured, the glochidia load was estimated from that one individual.

Juvenile survival rate was measured using a modified Buddensiek mesh cage (Buddensiek, 1995) with a 130 mm long axis, 90 mm short axis, 10 mm thickness, 5 mm cell diameter, and 250 μm mesh nets. The juveniles were collected by artificial infection using glochidia of six gravid mussels from site *u* and approximately 500 YOY white-spotted char individuals (see Appendix 6). On 29-31 July, 2018, five live juveniles were placed in each cell, giving a total of 30 individuals per cage. Triplicate mesh cages were installed on the riverbed in similar environments as much as possible in each reach (a total of 90 individuals/site) between 30 July and 1 August 2018. Surviving juveniles in each mesh cage were counted on three occasions: 27 August–1 September (1^st^ monitoring), 27–29 September (2^nd^ monitoring), and 22–24 October (3^rd^ monitoring). Mussel survival was judged by visible juvenile activities (valve movement and/or foot activities) using a stereomicroscope in the field.

The experiment was conducted in accordance with Hokkaido University legislation on animal experimentation (Hokkaido University, 2007).

### Environmental factors

Inorganic fractions of suspended substances (“fine sediment”, hereafter) were measured as an indicator of fine sediment abundance. Surface water samples (4 L) were collected from the streamflow at each site on 27–29 September 2018. The collected samples were taken to the laboratory for subsequent treatment. Water samples were filtered using pre-combusted and pre-weighed micro fibre glass filters (0.7 μm pore size). Fine sediment (mg/L) was measured through loss on ignition after drying for >24 h at 60 °C and combustion for 3 h at 450 °C.

Electrical conductivity (EC, mS/m) had a strong positive correlation with total nitrogen (TN) and total phosphorus (TP) in river surface water in the study region (*r* = 0.88 and 0.45, *P* <0.05, N = 24), suggesting that EC could be used as an indicator of nutrient concentration in the rivers (Ishiyama et al., 2020). EC was measured (WM-32EP, TOA DKK Co., Japan) at the uppermost part of each reach during monitoring of mussel gravidity, when fish sampling, and at each monitoring of the field experiment.

### Statistical analyses

We constructed generalised linear models (GLMs) to test the effects of three life-cycle stage indices on the proportion of juveniles as the response variable. In addition, the interaction of glochidia density and juvenile survival rate was included as an explanatory variable because they had exhibited a positive interaction with the proportion of juveniles in preliminary plots. For juvenile survival rate, the mean values at the three monitoring times were used as explanatory variables. The survival rate during the 2^nd^ and 3^rd^ monitoring at site *i* was excluded from further analyses because all experimental cages had become buried in the streambed by >30 cm, which was too deep for the juveniles. We developed all possible models by progressively removing each of the explanatory variables, and multiple models were compared based on Akaike’s information criterion (AIC) (Akaike 1974). We considered models with ΔAIC <2 to be meaningful representations of the relationship between variables (Burnham & Anderson, 2002). Model averaging was performed for models with ΔAIC <2. The explanatory parameters with 95% confidence intervals (CI) that did not include zero were considered influential parameters in each averaged model. When one model fell within ΔAIC <2, we tested the significance of each explanatory variable in this model using Wald Z statistics.

As a result of the above model selection, the interactive effect of glochidia density and juvenile survival rate on the proportion of juveniles was positive among the significant variables (see “Results” section). We constructed GLMs and generalised linear mixed models (GLMMs) to determine the limiting factors of glochidia density and juvenile survival rate, respectively. To explain glochidia density, we used gravid density, EC values measured on fish sampling occasions, fine sediment, and the interaction of EC and fine sediment. To determine the juvenile survival rate, fine sediments, EC averaged by values on three monitored juveniles, and the interaction of the two variables were used as explanatory variables. The timing of monitoring (1^st^, 2^nd^, or 3^rd^) was used as a random factor. Model selection and determination of influential parameters were conducted as described above.

In GLMs and GLMMs, binomial error distribution was performed using a log link function for ratio data as a response variable, whereas Gaussian error distribution was used for other data types. We confirmed that there was no significant multicollinearity among explanatory variables when constructing each model (r <0.60 with *P* >0.05, in all cases). All statistical analyses were conducted using R 3.0.3 (R Development Core Team, 2015) with a significance level of α = 0.05. The glmmADMB, lme4, and MuMIn packages were used.

## Results

### Population status

A total of 2,491 FPM individuals was collected. More than 50 individuals were caught (N = 55–211) at each site, except for four sites (sites *l, m, r*, and *s*, N = 14–47). The proportion of juveniles ranged between 0.00 and 0.53, and juveniles were absent from four sites (sites *p, q, s*, and *w*) (Fig. 2).

**Fig. 2.**
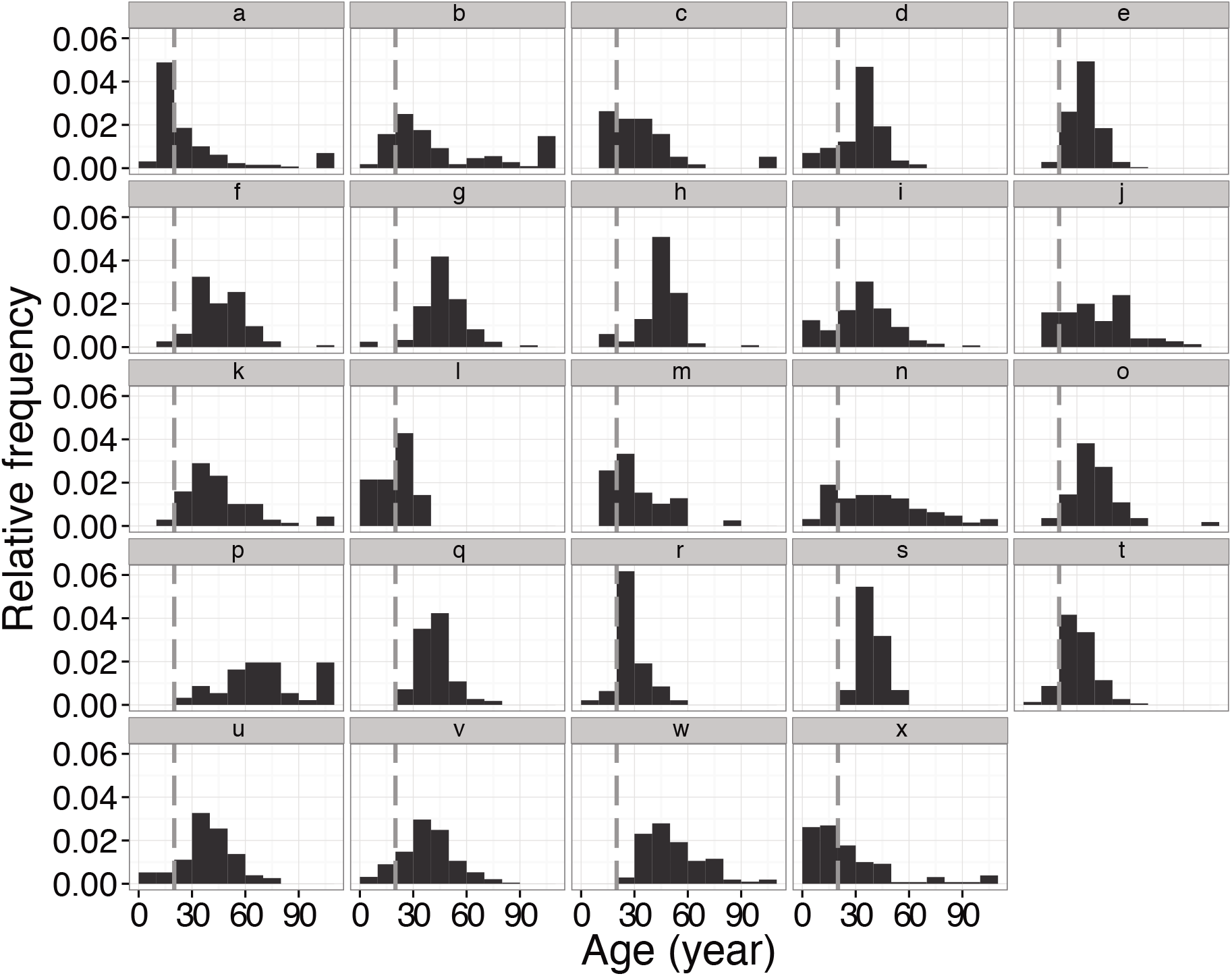
Relative frequency distributions of the age of *Margaritifera togakushiensis* populations in 24 study rivers (sites). All individuals aged >100 years were categorised in the same category. Grey dashed lines indicate the age of 20-year-old juveniles.

### Key life-cycle stage status

The gravidity of 1,260 FPM individuals was checked on two monitoring occasions. On each occasion, the number monitored at each site ranged between 31 and 81 (57.27 ± 11.61) individuals/site. The gravid density of each site ranged between 0.00 and 18.51 (6.77 ± 6.50) individuals/m^2^.

A total of 277 host individuals (90 YOY and 187 ≥1^+^ individuals) were collected. Infection of FPM glochidia was confirmed in the host samples from all sites. A total of 191 host individuals (48 and 143 individuals of YOY and ≥1^+^, respectively) were examined for glochidia load. The mean glochidia load of YOY individuals ranged between 15.00 and 134.00 (65.98 ± 41.29) among six sites and of ≥1^+^ individuals between 130.24 and 2045.31 (830.30 ± 753.88) among 11 sites (Appendix 1: Table S1). The glochidia density of each site ranged between 3.17 and 65.51 (29.81 ± 22.83) individuals/m^2^.

The juvenile survival rates ranged between 0.00 and 0.78 (0.39 ± 0.30), 0.00 and 0.74 (0.35 ± 0.27), and 0.00 and 0.64 (0.28 ± 0.24) on 1^st^, 2^nd^, and 3^rd^ monitoring, respectively. The juvenile survival rate decreased by an average of 0.61 for the first month, and this decreasing trend continued to the end (Appendix 1: Fig. S3).

The proportion of juveniles was modelled using three life-stage indices and the interaction of glochidia density and mean juvenile survival rate (Appendix 1: Table S2). Among the four explanatory variables, the effect of the interaction of glochidia density and mean juvenile survival rate was significantly positive (*P* <0.001) (Table 1). The positive effect of glochidia density on recruitment success was more significant in sites with higher juvenile survival rates (Fig. 3). The effect of the gravid density was also statistically significant (*P* <0.001) but negative (Table 1).

**Table 1.** Parameter estimates and *P*-values for the Wald-Z statistics of the best model explaining the proportion of juveniles.

| Explanatory variable | Estimate | SE | *P*-value |
| --- | --- | --- | --- |
| Gravid density | -0.13 | 0.02 | <0.001 |
| Glochidia density | -0.01 | 0.01 | 0.26 |
| Mean juvenile survival rate | -0.73 | 0.76 | 0.34 |
| Glochidia density ×<br>Mean juvenile survival rate | 0.09 | 0.02 | <0.001 |

**Fig. 3.**
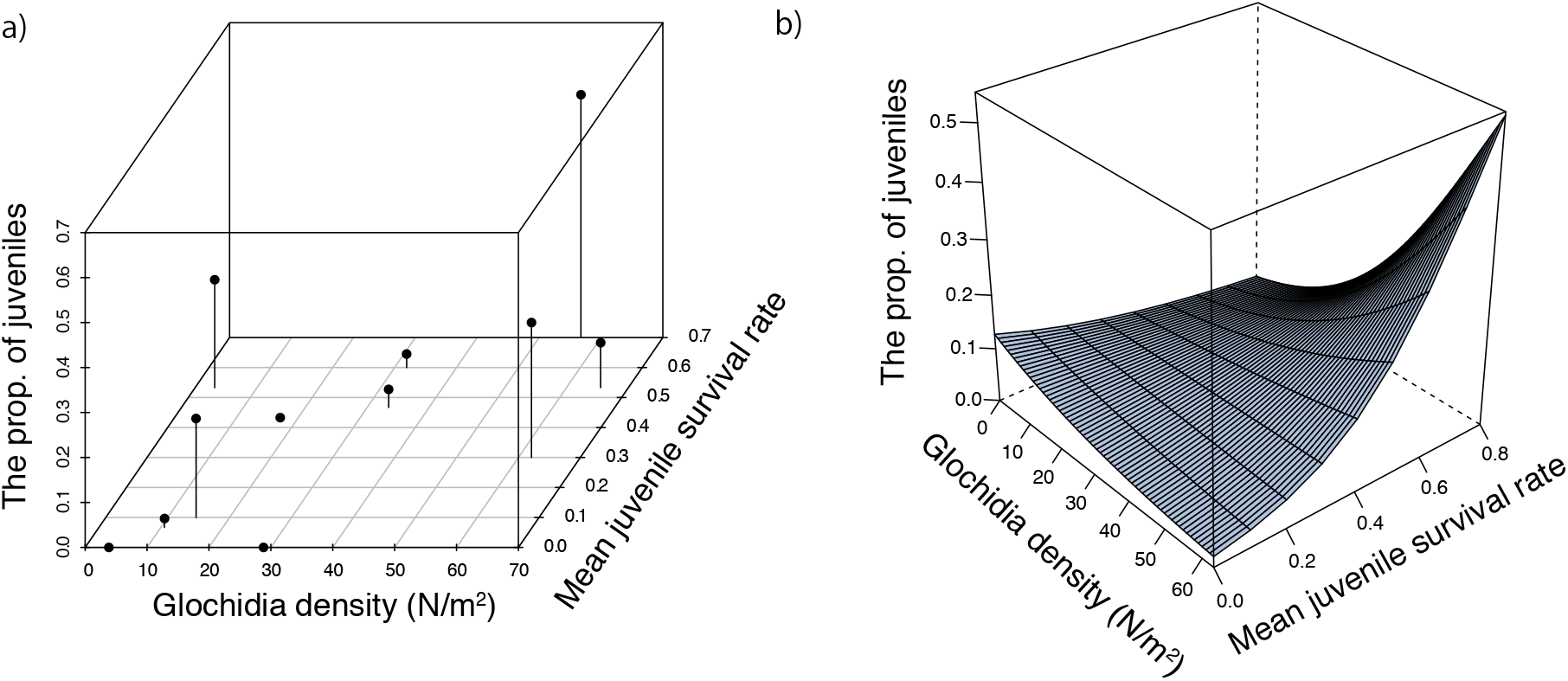
Interactive effects of mean juvenile survival rate and glochidia density (N/m^2^) on the proportion of juveniles. Actual measurements of the variables (a). Estimates of the proportion of juveniles using the generalised linear model (GLM) with a binomial error distribution (b).

The model selection results for explaining the glochidia density showed that the top eight models included EC, gravid density, and fine sediment (Appendix 1: Table S3). The 95% confidence intervals of all factors included zero, indicating non-meaningful effects (Table 2).

**Table 2.** Model averaged parameter estimates and 95% confidence intervals for the models with ΔAIC <2 that explain glochidia density.

| Explanatory variable | $\beta$ | SE | 95% confidence interval | |
| --- | --- | --- | --- | --- |
|  |  |  | lower | upper |
| Gravid density | 1.58 | 0.92 | -0.57 | 3.74 |
| Fine sediment | -4.02 | 2.69 | -10.36 | 2.33 |
| EC | -0.86 | 0.77 | -2.66 | 0.94 |

$\beta$ : Model averaged estimates of parameters

Among the models explaining the juvenile survival rate, the top one included fine sediment, EC, and their interaction as explanatory variables (Appendix 1: Table S4). Among the three explanatory variables, the effect of the interaction of EC with fine sediment was significantly negative (*P* <0.001) (Table 3). The negative effect of fine sediment became more apparent in sites with higher EC values (Fig. 4).

**Table 3.**
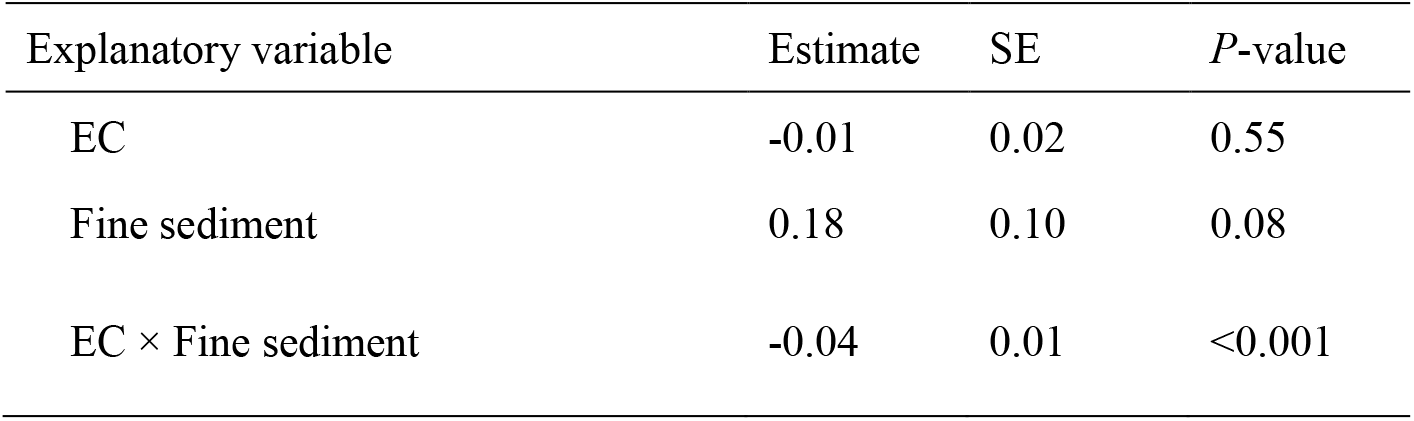
Parameter estimates and *P*-values for the Wald-Z statistics of the best model explaining the juvenile survival rate.

**Fig. 4.**
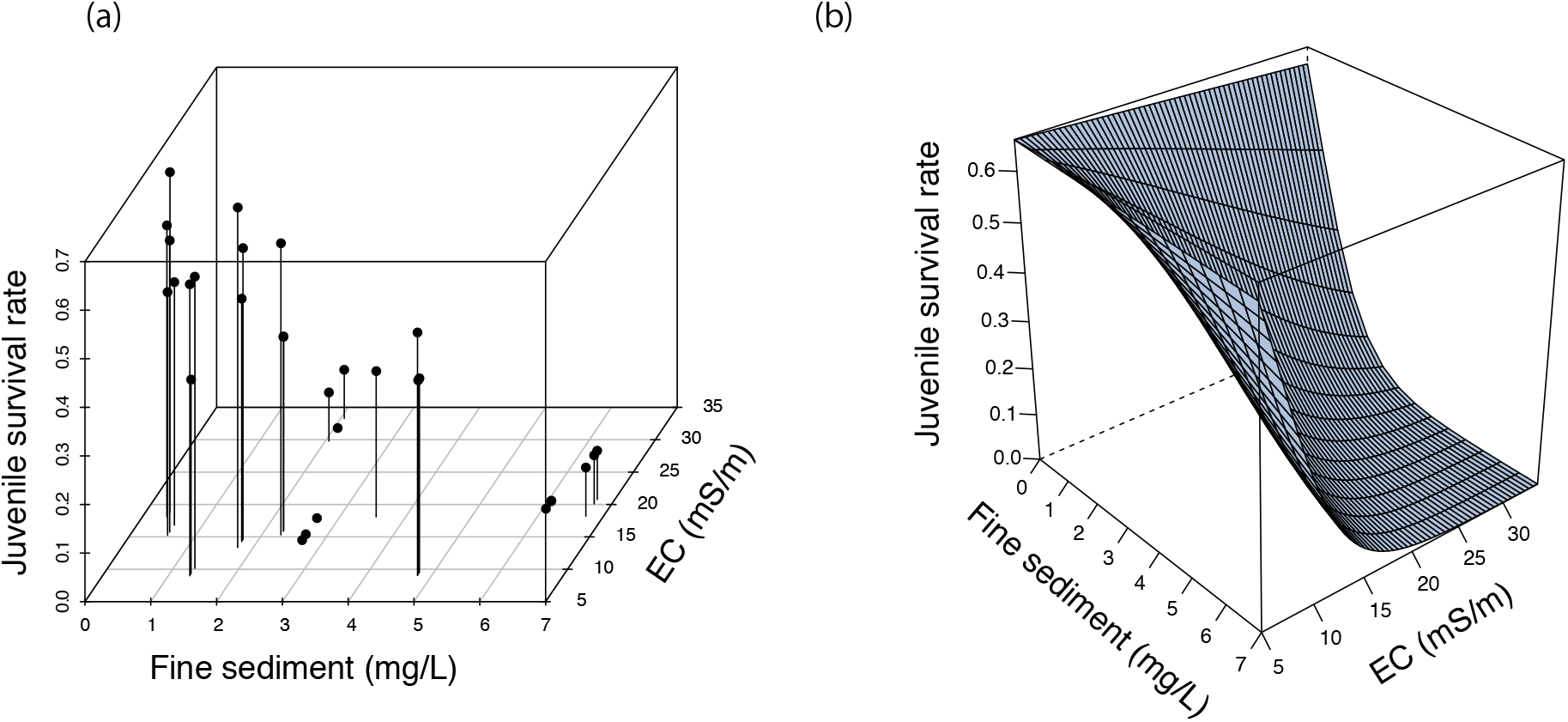
Interactive effects of electrical conductivity (EC, ms/m) and fine sediment (mg/L) on juvenile survival rate. Actual measurements of the variables (a). Estimates of juvenile survival rate using the generalised linear mixed model (GLMM) with a binomial error distribution (b).

## Discussion

Despite the known effects of multiple stressors on population responses by using abundance or presence/absence data of organisms (e.g., Toft et al., 2018), the effect of multiple stressors on recruitment efficiency remains unclear. This knowledge gap is particularly wide for organisms with complex life cycles. To the best of our knowledge, this is the first study to show that increases in nutrient concentration and fine sediments interactively cause recruitment failure of freshwater mussels by synergistically reducing the survival rate of early juveniles. In addition, the present study showed that a reduced potential supply of juveniles (i.e. low glochidia density) regulated at the parasitic stage could cause recruitment failure interactively with the juvenile survival rate. Gravid density regulated at the adult stage had a negative relationship with the proportion of juveniles for unclear reasons. However, it is unlikely that lower larval production in the population (i.e. lower gravid density) would cause higher reproductive efficiency (see “Theoretical framework”). Therefore, this stage could not be a bottleneck in recruitment efficiency. Recruitment failure is a major cause of the decline in freshwater mussels (Strayer & Malcom, 2012). Previous studies have suggested that habitat degradation from excessive nutrients and fine sediment causes recruitment failure of freshwater mussels via impacts on juvenile survival rate (Geist & Auerswald, 2007; Österling et al., 2010; Strayer & Malcom, 2012). However, they could not distinguish the effects of the two stressors on the post-parasitic juvenile stage and earlier life stages or test whether they interacted. Our key findings suggest a synergistic effect of multiple stressors on the performance of the post-parasitic juvenile stage, creating a critical bottleneck in the declining population of freshwater mussels, and highlighted the need for management of multiple stressors.

The negative effects of nutrients and fine sediment on mussels were consistent with the previously reported independent effects of each. Nutrients generally have no strong acute toxicity but can have weak or chronic toxicity to mussel juveniles. For example, high concentrations of nitrate (NO_3_^-^), which was a major component of the nutrients that had a strong positive relationship with EC at our sites (*r* = 0.86, *P* <0.01; N = 24 sites; unpublished data), can reduce the juvenile survival rate (Douda, 2010). In addition, excessive fine sediment in the water column can directly decrease the survival rate of juvenile mussels by clogging their gills (Österling et al., 2010). However, the reasons for the synergistic effect of the two stressors on juvenile survival are unclear. Possible mechanisms may involve modifications of the biogeochemistry in the benthic zone. Sustained high levels of fine sediment in water could cause sediment deposits on the riverbed and clogging of the interstitial spaces (Fung & Ackerman, 2019), which potentially disrupts water exchange at the surface-hyporheic interfaces. This, in turn, may cause hypoxic conditions and higher concentrations of un-ionised ammonia that are strongly toxic to juvenile mussels (Strayer & Malcom, 2012; Wood & Armitage, 1997). Furthermore, increasing nutrient concentrations may exacerbate this condition because of loading with reactive nitrogen as the source of ammonia becomes more abundant (Strayer & Malcom 2012).

Glochidia density was found to be important for the recruitment of FPM, together with the juvenile survival rate. The lack of juvenile supplies to the population may eventually decrease the number of surviving juveniles. On the other hand, glochidia density had a weak relationship with abiotic and biotic factors, suggesting that it is not strongly limited by these factors. The glochidia density is a function of host density and mean glochidia load per fish. Low habitat quality can reduce host fish availability and the infection success of glochidia larvae (Kawajiri et al., 2021). However, the host fish density in the study region was mainly limited by catchment size and habitat complexity and not by habitat degradation (Ishiyama et al., 2020). In addition, the relationships between glochidia load of both age classes (YOY and ≥1^+^) and fine sediment and nutrient concentrations were also unclear (GLM with log-likelihood ratio tests, YOY: *P* = 0.92 and *P* = 0.38; ≥1^+^: *P* = 0.24, and *P* = 0.62). Therefore, the indirect effects of habitat degradation in the study region on mussel recruitment of FPM through its effects on host population size were considered small.

We found that the lack of glochidia density regulated at the parasitic stage can also cause recruitment failure interactively with the juvenile survival rate; the positive effect of glochidia density on the reproduction success of FPM became more significant in sites with higher juvenile survival rates. This finding also has meaningful management implications, namely that both parasitic and post-parasitic stages should be considered when re-establishing recruitment of freshwater mussels. Previous studies attempting to re-establish mussel recruitment have not always been successful. For example, Galbraith et al. (2018) reported that the reintroduction of American eels *Anguilla rostrata*, which is a migratory host fish of *Elliptio complanata*, could re-establish recruitment of *E. complanata* at only one of two target sites where American eels were absent due to fragmentation by past dam installation. This discrepancy in the results between the two sites might be caused, at least partially, by the decreased juvenile survival rate due to habitat degradation. Therefore, enhancing glochidia density by improving habitat quality is also important for increasing the natural recruitment level of freshwater mussels.

Given the significant interaction between nutrients and fine sediment, several management options can be proposed to overcome the bottleneck of low survival rates of early juveniles. The positive effects on the juvenile survival rate of reduced nutrient loading could be maximised in rivers carrying quantities of fine sediment. Both nutrients and fine sediments are generally related to land use, such as agricultural land cover in catchments (Howarth et al., 2012; Wood & Armitage, 1997). In the present study, EC had strong positive relationships with agricultural and urban land cover in the catchments (Appendix 1: Fig. S4, *r* = 0.73 and 0.86 with *P* <0.01, N = 24 sites), indicating that nutrients originated from agricultural land and urban areas. Restoring wetlands and vegetated buffer strips may effectively capture nutrients before they reach streams (Weller et al. 2011). In addition, increasing the percentage of households with sewerage connections may be important to reduce nutrient inputs into rivers from urbanised areas. Furthermore, establishing drainage and wastewater treatment plants or septic tanks can effectively reduce the pollution effects of domestic and industrial wastewater. Regarding fine sediment abundance, we found no clear relationship with agricultural land cover in the catchments (*r* = 0.02, *P* = 0.92, N = 24 sites). This might be explained by fine sediment coming from point sources rather than diffuse sources. Large amounts of fine sediments can enter river water after localised events in the catchment, such as landslides, riverbank erosion, and deforestation (Wood & Armitage, 1997). Determining the sources of fine sediment is an important future challenge, but with current uncertainty about their anthropogenic origins, management actions are prioritised towards controlling nutrients.

Our results suggest that enhancing glochidia density can also be important for maximising the positive effects of mitigating stressors. In particular, managing these two stressors can be most effective in streams with higher glochidia density but a low juvenile survival rate. Glochidia density can be increased by enhancing the biological control of both host density and glochidia load on the host fish. To increase host density, installing large wood structures in the rivers may be effective because they are a major determinant of fish habitat quantity and quality in general (He et al. 2009). The installation of these structures can enhance fish density by providing cover habitats and enhancing habitat heterogeneities, such as greater variations in water depth, current velocity, and the size of riverbed sediments (Nagayama et al. 2009). In addition, the (re-)introduction of host fish can increase host fish density. In particular, (re-)introduction during the reproductive period of FPM can be a more effective way to increase host fish density and the glochidia load on host fish. Furthermore, artificial infection of host fish with glochidia larvae may also be a more practical way to increase the glochidia load on individual fish because a larger number of glochidia can be attached more reliably (Preston et al. 2007). This artificial propagation might also be an important conservation measure for recovering the recruitment of FPM in the future.

## Supporting information

Appendix1_SuppInfo

Appendix2_SuppInfo

Appendix3_SuppInfo

Appendix4_SuppInfo

Appendix5_SuppInfo

Appendix6_SuppInfo

## Author’s contribution

KM, NI, JNN, DI, KK, TI, MN, FN conceived the ideas and designed the methodology; KM, NI, DI, TI, and HI collected the data; KM, NI, and JNN processed and analyzed the data; and KM led the writing of the manuscript. All authors contributed critically to the drafts and gave final approval for publication.

## Acknowledgments

We are grateful to Dr. Kishida of the Tomakomai Experimental Forest, Hokkaido University for providing an aquarium facility. We also thank members of Akkeshi Marine Station, Hokkaido University for helping with field logistics. We would like to thank Editage (www.editage.com) for English language editing. This study was partly supported by the Takara Harmonist Fund, Pro Natura Fund, Grand-in-Aid for Science Research of Lake Akkeshi and Bekanbeushi Wetland, JSPS Research Fellow Grant (JP18J12458), and the Environment Research and Technology Development Fund [S15 Predicting and Assessing Natural Capital and Ecosystem Services (PANCES)] of the Environmental Restoration and Conservation Agency, Japan.

