## Appendix1_SuppInfo for "Multiple stressors and recruitment failure of long-lived endangered freshwater mussels with a complex life cycle"

### Supplemental tables

**Table S1** The summary of the number of fish samples and mean glochidia load of two age classes (YOY and $\geq 1^{+}$ individuals) in the 11 study rivers (sites).

| River | The number of collected host fish [total number (YOY)] | The number of dissected host fish [total number (YOY)] | Mean glochidia load (YOY) | Mean glochidia load ($\geq 1^{+}$) |
|---|---|---|---|---|
| *c* | 11 (0) | 11 (0) | - | 384.00 |
| *d* | 10 (8) | 10 (8) | 80.88 | 1689.50 |
| *e* | 24 (0) | 14 (0) | - | 1282.79 |
| *f* | 35 (17) | 35 (17) | 40.88 | 131.47 |
| *i* | 17 (0) | 13 (0) | - | 2045.31 |
| *n* | 17 (0) | 17 (0) | - | 130.24 |
| *p* | 15 (1) | 10 (1) | 15.00 | 534.22 |
| *q* | 24 (1) | 18 (1) | 48.00 | 302.94 |
| *u* | 82 (64) | 36 (20) | 77.15 | 173.13 |
| *w* | 15 (0) | 15 (0) | - | 533.60 |
| *x* | 28 (1) | 15(1) | 134.00 | 1926.14 |

YOY: Young of the year

**Table S2** Summary of the model selection results relating variables of three life-cycle stage indices to the proportion of juveniles.

| Variables | df | AIC | ΔAIC | Weight |
|---|---|---|---|---|
| Grav + Glochi + Surv + Glochi × Surv | 5 | 160.79 | 0.00 | 1.00 |
| Grav + Glochi + Surv | 4 | 179.73 | 18.93 | 0.00 |
| Grav + Glochi | 3 | 205.78 | 44.99 | 0.00 |
| Grav + Surv | 3 | 208.22 | 47.43 | 0.00 |
| Glochi + Surv + Glochi × Surv | 4 | 230.73 | 69.94 | 0.00 |
| Glochi + Surv | 3 | 258.74 | 97.94 | 0.00 |
| Glochi | 2 | 267.64 | 106.84 | 0.00 |
| Surv | 2 | 268.98 | 108.18 | 0.00 |
| Grav | 2 | 275.97 | 115.18 | 0.00 |
| null | 1 | 302.37 | 141.58 | 0.00 |

"Grav"=Gravid density, "Glochi"= Glochidia density, "Surv"=Juvenile survival rate

**Table S3** Summary of the model selection results relating gravid density, EC, and fine sediment to the glochidia density.

| Variables | df | AIC | ΔAIC | Weight |
|---|---|---|---|---|
| Grav | 3 | 101.75 | 0.00 | 0.18 |
| Fine + Grav | 4 | 101.90 | 0.15 | 0.17 |
| EC + Fine | 4 | 102.82 | 1.07 | 0.11 |
| null | 2 | 102.98 | 1.23 | 0.10 |
| EC + Fine + Grav | 5 | 103.00 | 1.25 | 0.10 |
| Fine | 3 | 103.08 | 1.33 | 0.09 |
| EC + Grav | 4 | 103.22 | 1.47 | 0.09 |
| EC | 3 | 103.28 | 1.53 | 0.08 |
| EC + Fine + Grav + EC × Fine | 6 | 104.20 | 2.45 | 0.05 |
| EC + Fine + EC × Fine | 5 | 104.76 | 3.01 | 0.04 |

"Grav"=Gravid density, "Fine"= Fine sediment

**Table S4** Summary of the model selection results relating EC, fine sediment, and interaction of the two variables to the juvenile survival rate.

| Variables | df | AIC | ΔAIC | Weight |
|---|---|---|---|---|
| EC + Fine + EC × Fine | 4 | 516.65 | 0.00 | 1.00 |
| EC + Fine | 3 | 556.97 | 40.31 | 0.00 |
| Fine | 2 | 769.74 | 253.09 | 0.00 |
| EC | 2 | 922.81 | 406.16 | 0.00 |
| null | 1 | 1168.87 | 652.22 | 0.00 |

"Fine"=Fine sediment

### Supplemental figures

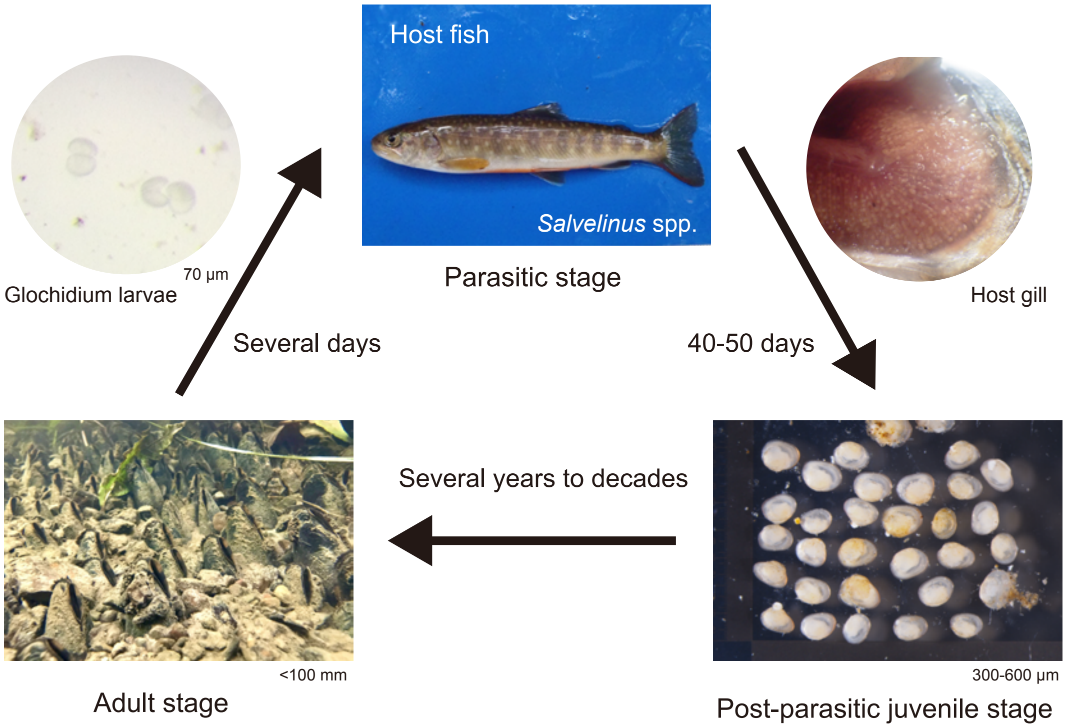


**Fig. S1** The general life cycle of *Margaritifera togakushiensis* as an example of Unionoid's life cycle.

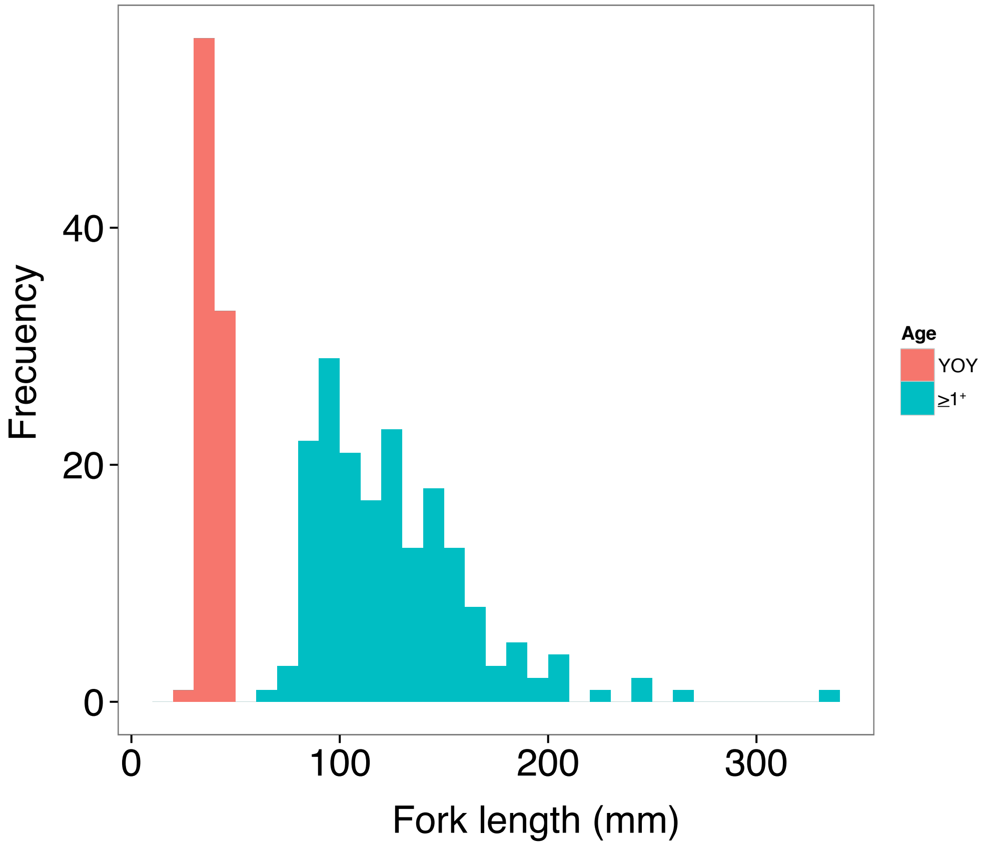


**Fig. S2** The histogram of fork length (mm) of host fish (White-spotted char *Salvelinus leucomaenis leucomaenis* and three ≥$1^{+}$ individuals of Dolly Varden char *S. malma krascheninnikovi*). Red and blue indicate YOY (young-of-the-year) and ≥$1^{+}$ individuals, respectively.

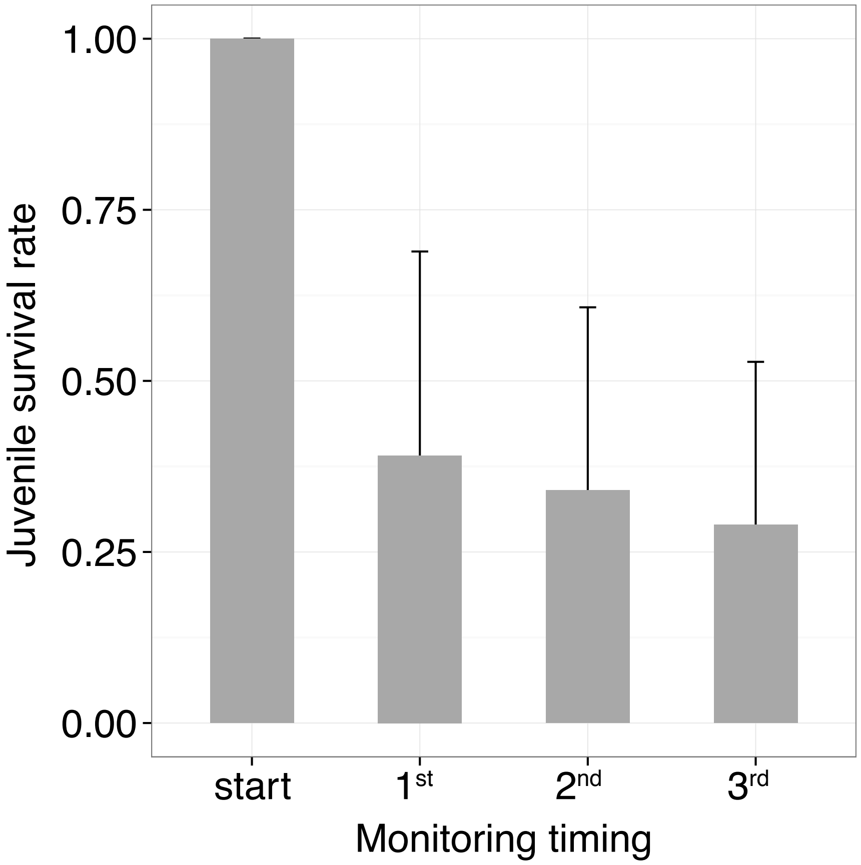


**Fig. S3** The bar plots of time changes in the juvenile survival rate of *Margaritifera togakushiensis*. The “start” means the beginning timing of field exposure experiments of juveniles. Error bars show standard deviations.

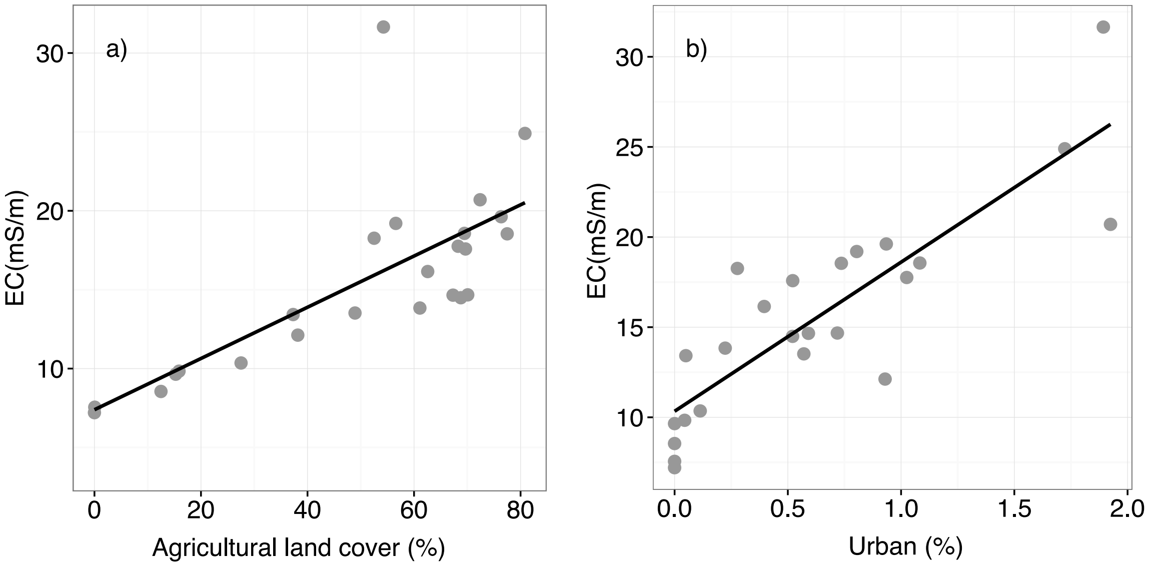


**Fig. S4** Relationships between electrical conductivity (EC, mS/m), and agricultural land (a) and urban (b) covers (%). The black lines were drawn using the regression in the linear models. The land use data were derived and calculated the protocols below (see "GIS analysis)".

GIS analysis.— The study derived watershed areas for the most downstream study reach in each study river using the Spatial Analyst Toolkit in ArcMap 10.6 (ESRI inc., Redland. CA, USA) with a 50-m digital elevation model (DEM) from the Geospatial Information Authority of Japan. We obtained land cover (100-m resolution) for the study region from the nation-wide land use census data (land use database in 1987, Ministry of Land, Infrastructure, Transport and Tourism). The present land cover in the study region was developed by the late 1980s (Nagasaka, 2017). The land use database in 1987 has mostly no change in the current land cover.

<u>References</u>


Nagasaka, A. (2017). *The restoration of Furen watersheds*. Sapporo: Hokkaido University Press.
