## Appendix2_SuppInfo for "Multiple stressors and recruitment failure of long-lived endangered freshwater mussels with a complex life cycle"

### **The protocols to assess the population status of *Margaritifera togakushiensis***

For each study reach, five transects were laid out at equal intervals. Three 0.09-m$^2$ quadrats were placed, one at mid-channel and two at both sides separately, near the bank, for a total of 15 quadrats per reach in each transect. Mussels were collected from each quadrat as follows. First, all visible mussels were collected using a glass-bottomed viewing bucket. Second, we excavated the bed-sediment to a depth of ~10-cm using a trowel and immediately sieved all material through a 2-mm mesh sieve to reduce the size bias of mussels (Strayer & Malcom, 2012). We collected at least 100 *Margaritifera* individuals from one study site to reach at least 50 individuals of *M. togakushiensis* except for site *l*. Only a total of fifty-seven *Margaritifera* individuals were collected from two study reaches of site *l* because the previous study revealed the distribution of the only *M. togakushiensis* in the site (Miura et al., 2019). When the number of collected *Margaritifera* individuals by quadrat sampling was less than 100 individuals in a study site, we additionally collected the mussel individuals to be a total of 100 or more individuals. Then, one investigator found visually mussel individuals using a glass-bottomed viewing bucket by random walking in the study reach, and we laid out a quadrat with found individuals as the center and collected mussels in the same manner as above procedures.

10.1890/11-1536.1
