## Appendix3_SuppInfo for "Multiple stressors and recruitment failure of long-lived endangered freshwater mussels with a complex life cycle"

### **The protocols of age determination and the results of fitting growth models for *Margaritifera togakushiensis***

Growth models of *M. togakushiensis* were developed to determine the age structure of the population and identify the ages of <20 years individuals. The age of margaritiferids can be estimated by the relationship between shell length and the number of growth rings on the ligament (e.g. Hastie et al. 2000). Age determination and development of growth models were performed using the protocol of Akiyama & Iwakuma (2010) based on collected 23–83 (mean: 42) live mussels from each watershed including the study sites from 2017 to 2018. The collected individuals were identified using the simple non-lethal criteria and/or the presence of mature glochidium larvae in the gills of female individuals on the occasion of the reproductive season of *M. togakushiensis* (Kondo, 2008; Miura et al., 2019). After the shell and ligament length of each individual were measured, the number of growth rings was counted under a stereomicroscope. A part of the ligaments of margaritiferids is usually abraded due to erosion. Thus, the number of growth rings on the abraded part of the shell was estimated on the basis of the relationship between shell length and the number of growth rings obtained from 28 non-eroded juvenile shells (shell length: 7.6–22.0 mm) collected from the study region done as with Hendelberg (1961). This study fitted four non-linear models: Bertalanffy growth function, Hyperbolic saturation function, Logistic function, and Gompertz function, to individuals of each watershed using measured shell length and age of individual. These models have been traditionally used to describe the growth characteristics for freshwater and marine bivalves including margaritiferids (Akiyama & Iwakuma, 2010; Haag & Rypel, 2011; Nakamura et al., 2018; San Miguel et al., 2004). The four models are expressed as the following functions.

Bertalanffy growth function

$$L_t = L_\infty(1 - e^{-k(t-t_0)})$$

Hyperbolic saturation function

$$L_t = \frac{L_\infty k(t - t_0)}{1 + k(t - t_0)}$$

Logistic function

$$L_t = \frac{L_\infty}{1 + e^{-k(t-b)}}$$

Gompertz function

$$L_t = L_\infty e^{-ae^{-kt}}$$

where $L_\infty$ is the theoretical maximum length (or asymptotic length, mm), $k$ is the growth coefficient ($year^{-1}$), $t_0$ is the theoretical age at zero-length (year), and $a$ and $b$ (year) are constants. The best-fitting model was selected by comparing RSS (residual sum of squares) (Akiyama & Iwakuma, 2010).

As a result of the comparisons of fitting curves of age estimation functions, Hyperbolic saturation function and Gompertz function were the most fitted function in seven study watersheds (*A, B, C, D, E, G,* and *H*) and one watershed (*F*) respectively, that is supported by the lowest RSS of the comparison of four functions in each watershed (Table A). The shell length of individuals estimated as 20-years in each watershed was 37.9–54.2 mm calculated from the length-age relationship (Table A).

**Table A** The summary of the results of fitting four non-linear models to *Margaritifera togakushiensis* individuals in each watershed.

| Watershed | River (site) | Most fitted function | RSS* | N** | Length (mm) of 20 yr individuals |
|---|---|---|---|---|---|
| *A* | *a, b, c* | Hyperbolic | 809.9 | 48 | 54.2 |
| *B* | *d* | Hyperbolic | 637 | 30 | 39.4 |
| *C* | *e* | Hyperbolic | 215.4 | 23 | 41.5 |
| *D* | *f, g, h* | Hyperbolic | 1072 | 42 | 37.9 |
| *E* | *i, j* | Hyperbolic | 745.1 | 46 | 47.3 |
| *F* | *k, l* | Gompertz | 306.6 | 29 | 39 |
| *G* | *m, n, o* | Hyperbolic | 856.1 | 31 | 41.6 |
| *H* | *p, q, r, s, t, u, v, w, x* | Hyperbolic | 1913 | 83 | 44.5 |

*RSS: residual sum of squares

**N: the number of samples
