## Appendix4_SuppInfo for "Multiple stressors and recruitment failure of long-lived endangered freshwater mussels with a complex life cycle"

One investigator randomly collected visible mussels using a glass-bottom viewing bucket for 15–30 min in each study reach. Shell length of 40 mm was considered as threshold size above while individuals of target species were sexually mature in the study region (Miura, 2020). Thus, at least 50 *M. togakushiensis* individuals with a more than 40-mm shell length were collected from each site. Each individual was checked for the presence of mature glochidia in the gills by slightly opening the shell valves (<5 mm) with a shell opener. The mussels with swollen and cream-colored outer demibranchs were considered to be gravid individuals (Hastie and Young 2003). Each mussel was photographed on the white tray with a ruler and a level gauge in relation to the gravidity condition (presence/absence of matured glochidia). Each collected mussel was marked with a small sign on the shell surface using white paint or a hand drill (Miura et al., 2018; Negishi & Kayaba, 2009), and each was immediately released to the original study reach after marking. If the marked mussels were captured on the occasion of later observation, those were excluded from targets of observation to avoid handling effects. Later, the shell length and height of each individual were measured from the photographed images as the same protocols in the "population status assessment" section in the main text, and *M. togakushiensis* individuals were identified by using the criteria of Miura et al. (2019).

The gravidity rate of *M. togakushiensis* at the first monitoring (0.31 ± 0.17) (mean ± SD) was higher than the second (0.14 ± 0.15) (Fig. A). The lower gravidity rate on 2$^{nd}$ monitoring means that the glochidia release had already occurred and thus measured values did not necessarily represent true gravidity rate. Therefore, the values of the gravidity rate of the first monitoring were used for the calculation of gravid density.

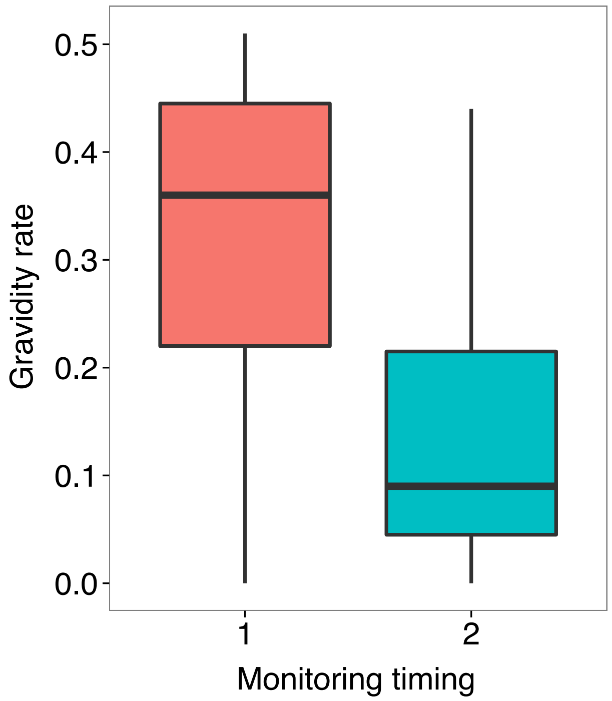


**Fig. A** Boxplots showing gravidity rate of *Margaritifera togakushiensis* on first and second monitoring occasions. The lower ends, top ends, and central thick lines of boxes represent 25% and 75% quartile ranges, and medians, respectively. Error bars indicate a maximum value within 1.5 times of the box height.
