## Appendix5_SuppInfo for "Multiple stressors and recruitment failure of long-lived endangered freshwater mussels with a complex life cycle"

### **The protocols to determine the host density and mean glochidia load**

The host fish (white-spotted char *Salvelinus leucomaenis leucomaenis* and Dolly Varden char *S. malma krascheninnikovi*) density (N/m$^2$) was estimated by the 2-pass electrofishing method (quantitative electrofishing) in reach with the lengths of 10 times of mean wetted-width in each study river. The fork length (mm) of each captured fish was measured after anesthetizing (FA100, DS Pharma Animal Health Co., Osaka, Japan). When the number of captured individuals of each age class [Young-of-the-year (YOY) or ≥$1^+$ individuals] in each study reach didn't reach 10 individuals at quantitative electrofishing, additional electrofishing was performed 1–2 h to qualitatively collect host fish individuals within the fish collection reach and 50–100 m upstream and/or downstream of the reach. To calculate the collection area, five transects were laid at the fish collection reach and wetted-width was measured on each transect using a 50-m tape. The host density (N/m$^2$) of YOY and ≥$1^+$ individuals were calculated respectively using the data of quantitative electrofishing.

Of the collected host fish individuals, more than 10 individuals of each age class were brought back to the laboratory and were frozen after anesthetizing. But only two rivers had more than ten samples of YOY individuals (Appendix 1: Table S1). In the laboratory, all gills (a total of eight-gill arches/fish) were removed from each individual, and the total number of glochidia infection on each fish was counted under a stereomicroscope in the laboratory.
