## Appendix6_SuppInfo for "Multiple stressors and recruitment failure of long-lived endangered freshwater mussels with a complex life cycle"

**The maintenance of aquaria for infected fish and procedures of juvenile production**

The juveniles of *Margaritifera togakushiensis* were collected by artificial infection using about 500 YOY white-spotted char individuals [fork length (FL): 25.00–40.00 mm]. These host fish individuals were produced by artificial insemination and cultured in indoor aquaria of Tomakomai Experimental Forest of Hokkaido University (Fig. B, lat. 42°40'N, long. 141°35'E). The origins of YOY individuals were six gravid females [FL: 462.00–668.00 (532.83 ± 90.83) (mean ± SD) mm] and 14 adult males [FL: 158.00–460.00 (224.43 ±71.97) mm] collected from the two watersheds including sites *e* and *p* to *x* in autumn 2017 (Ito, 2020). Six gravid mussels [shell length: 50.00–68.00 (59.58 ± 6.71)] mm were collected from site *u* for the infection on 18 May 2018. The host fish was exposed to a well-aerated glochidia slurry (40,000 N/L) derived from the six gravid females for 30 minutes on 19 May 2018 and kept until the excystment of juveniles. More detailed infection procedures are shown in Ito (2020). The peak of juvenile excystment occurred from 23–28 July 2018 and individuals were collected gently using Pasteur pipettes under a stereomicroscope. Only juveniles collected during the peak period were used for the experiments to avoid bias by using incompletely developed juveniles.

**Fig. B** Geographical location of the Tomakomai Experimental Forest of Hokkaido University (a), aquaria in the experimental forest (b), and infected YOY individuals of white-spotted char in the aquarium (c).

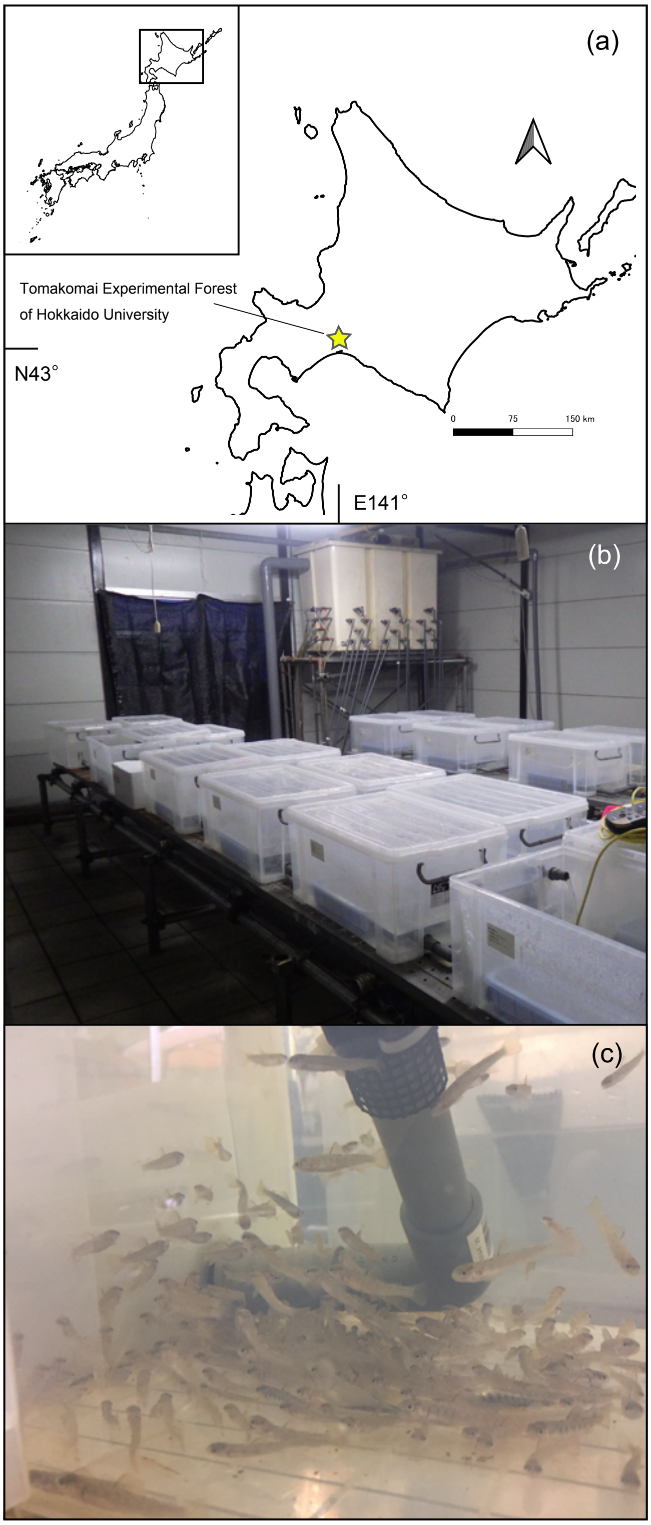
